# Model Adequacy Tests for Likelihood Models of Chromosome-Number Evolution

**DOI:** 10.1101/2020.10.05.326231

**Authors:** Anna Rice, Itay Mayrose

## Abstract

- Chromosome number is a central feature of eukaryote genomes. Deciphering patterns of chromosome-number change along a phylogeny is central to the inference of whole genome duplications and ancestral chromosome numbers. ChromEvol is a probabilistic inference tool that allows the evaluation of several models of chromosome-number evolution and their fit to the data. However, fitting a model does not necessarily mean that the model describes the empirical data adequately. This vulnerability may lead to incorrect conclusions when model assumptions are not met by real data.
- Here, we present a model adequacy test for likelihood models of chromosome-number evolution. The procedure allows to determine whether the model can generate data with similar characteristics as those found in the observed ones.
- We demonstrate that using inadequate models can lead to inflated errors in several inference tasks. Applying the developed method to 200 angiosperm genera, we find that in many of these, the best-fitted model provides poor fit to the data. The inadequacy rate increases in large clades or in those in which hybridizations are present.
- The developed model adequacy test can help researchers to identify phylogenies whose underlying evolutionary patterns deviate substantially from current modelling assumptions and should guide future methods developments.

## Introduction

Chromosome number is widely recognized as a key feature of eukaryote genomes. Its popularity in cyto-taxonomical and evolutionary studies has been attributed to its ability to provide a concise description of the karyotype, the ease by which it can be recorded, and its stable phenotype across repeated measurements. Processes that lead to changes in chromosome numbers have direct consequences on central evolutionary processes related to reproductive isolation and speciation, thus providing important information for species determination and phylogenetic relationships (Guerra, 2008; Weiss-Schneeweiss & Schneeweiss, 2013). While chromosome numbers generally exhibit strong phylogenetic signal (e.g. Vershinina & Lukhtanov, 2017; Carta *et al*., 2018), they are also highly dynamic. This variability has been particularly well acknowledged in plants, with counts ranging from *n* = 2 to *n* = 720 (Khandelwal, 1990; Ruffini Castiglione & Cremonini, 2012), and records showing intraspecific variation in 23% of angiosperm species (Rice *et al*., 2015). Understanding the underlying processes that gave rise to these changes allows inference of major genomic events that have occurred in the history of a clade of interest and the processes that have shaped its diversification.

Of the various mechanisms underlying chromosome-number change, polyploidy, or whole genome duplication (WGD) has received significant attention because of the profound impacts such an event has on the organism. Polyploids often differ markedly from their progenitors in morphological, physiological, or life history characteristics, which may contribute to their establishment in novel ecological settings (Stebbins, 1971; Levin, 1983; Ramsey & Schemske, 2002; Soltis *et al*., 2007; Leitch & Leitch, 2008; Ramsey & Ramsey, 2014; Spoelhof *et al*., 2017; Rice *et al*., 2019). Polyploidy is thus recognized as one of the major processes that has driven and shaped the evolution of higher organisms. A more subtle change in chromosome number is dysploidy, leading to step-wise changes in the number of chromosomes, but typically does not immediately alter the genomic content. Dysploidy occurs via several types of genome rearrangements, leading to ascending or descending dysploidy through chromosome fission or fusion (Weiss-Schneeweiss & Schneeweiss, 2013). Deciphering the pattern of chromosome-number change within a clade allows inferring the number and type of transitions that have occurred along branches of a phylogeny, to estimate ancestral chromosome numbers, and to categorize extant species as diploids or polyploids.

In the last decade, several tools that infer changes in chromosome numbers along a phylogeny were developed (Mayrose *et al*., 2010; Hallinan & Lindberg, 2011; Glick & Mayrose, 2014; Freyman & Höhna, 2017; Zenil-Ferguson *et al*., 2017, 2018; Blackmon *et al*., 2019). Among these, the chromEvol probabilistic framework (Mayrose *et al*., 2010) was the first to incorporate a continuous time Markov process that describes the instantaneous rate of change from a genome with *i* haploid chromosomes to a genome with *j* haploid chromosomes via specific types of dysploidy and polyploidy transitions. Further development of this framework allowed for more intricate types of chromosome-number transitions (Glick & Mayrose, 2014), to differentiate between transitions that coincide with speciation events and those that occur continuously in time along branches of the phylogeny (Freyman & Höhna, 2017), and to associate patterns of chromosome-number change with the evolution of a discrete character trait (Zenil-Ferguson *et al*., 2017; Blackmon *et al*., 2019).

In the chromEvol model, each type of transition is represented by a parameter describing its rate of change. The inclusion (or exclusion) of different parameters entails different hypotheses regarding the pathways by which the evolution of chromosome number proceeded in the clade under study. In a regular application of the chromEvol framework, different models are fitted to the data and the best one is chosen by comparing the relative fit of each model to the data at hand using established model selection criteria, such as the likelihood ratio test or Akaike Information Criterion (AIC; Akaike, 1974). In reality, however, no empirical dataset will meet all the assumptions of any model and thus relaying on the best model (or set of models) may be vulnerable to incorrect conclusions in datasets whose underlying evolutionary process deviate substantially from current modelling assumptions. To prevent such errors, here we develop a model adequacy test that allows determining whether a given model of chromosome-number evolution provides a realistic description of the evolutionary process for reliable inferences.

Several assumptions made by existing models of chromosome-number evolution may be violated when empirical data are analyzed. For example, all models rely on a memory-less Markovian process, in which the transition rates are only dictated by the current number of chromosomes of the lineage. Thus, for example, the transition rate from *n* = 10 to *n* = 9 is not affected by the duration of time the lineage possessed 10 chromosomes, nor by the sequence of events that had led to it. However, because rates of descending dysploidy may increase following WGD (Wood *et al*., 2009; Wendel, 2015; Soltis *et al*., 2016), the transition from *n* = 10 to *n* = 9 is more probable if *n* = 5 was the ancestral state compared to *n* = 11. Additionally, most models assume that the transition rates are similar across the phylogeny, although in practice the transition patterns may be rather different in some sub-clades compared to others, as has been demonstrated, for example, in Cyperaceae (Márquez-Corro *et al*., 2019). Finally, all current models are based on a phylogenetic structure and thus ignore the possibility of hybridizations. Notably, allopolyploidy, one of the main types of polyploidy, is defined by such reticulate evolutionary events and the biases caused by their presence is rather unexplored.

One aspect of understanding the reliability of a model and interpreting its results is to quantify its adequacy for the data and the question at hand. The aim of model adequacy tests is to determine the absolute fit of a model to the data, rather than to compare its relative fit among a set of models. With some variations, the general procedure of such tests is composed of several steps: first, given an empirical dataset, obtain the best-fitting model and its parameter values. Next, use that model to generate multiple simulated datasets. Then, compute several test statistics that describe various characteristics of the data on each simulated dataset and on the empirical dataset. If the empirical values of the test statistics fall outside the range of variation encompassed by the simulated data, then it may be concluded that the model cannot provide an adequate description of the data at hand. To date, model adequacy approaches are established for several types of data and inference tasks, including those related to sequence evolution (Bollback, 2002; Brown, 2014; Duchêne *et al*., 2015; Chen *et al*., 2019) and continuous valued organismal traits (Slater & Pennell, 2013; Pennell *et al*., 2015). However, both are inappropriate for data and analyses concerning the evolution of chromosome numbers as the former rely on statistics derived from many sites, while the latter rely on Brownian motion statistics.

In the following, we first provide the details of the developed model adequacy framework for likelihood models of chromosome-number evolution. We then use simulations to assess the type I error rate and to explore the consequences of using inadequate models in several common inference tasks, such as ancestral reconstructions of chromosome numbers and ploidy-level inference. Finally, we apply the developed procedure to a large cohort of angiosperm genera, as well as to clades that are expected to violate model assumptions.

## Methodological Description

### Model adequacy framework for chromosome-number evolution

Given chromosome count data and a compatible phylogeny (together denoted as *D*), chromEvol can be used to assess the fit of various models (*M*_*1*_, *M*_*2*_, …, *M*_*N*_ ; *N* denotes for the number of models) to *D*. Each model differs with respect to the included rate parameters or the constraints placed on them [θ(*M*_*1*_), θ(*M*_*2*_), …, θ(*M*_*N*_)]. The most general model considered here includes six free parameters (Glick & Mayrose, 2014) and assumes that five types of events are possible: a single chromosome-number increase (ascending dysploidy with rate *λ*) or decrease (descending dysploidy with rate *δ*), WGD (i.e. exact duplication of the number of chromosomes with rate *ρ*), demi-polyploidy (multiplications of the number of chromosomes by 1.5 with rate *μ*), and base-number transitions (the addition to the genome by any multiplication of an inferred base number, where *β*, is the inferred base number and *ν* is its respective transition rate). A combination of these parameters allows a range of models to be evaluated (Table 1 shows the various models considered here). We note that the chromEvol software also allows the ascending and descending dysploidy rates to depend on the current number of chromosomes, but this option was not evaluated here.

**Table 1.** The set of chromEvol models examined in this study, together with their rate parameters.

| Model | Model parameters<sup>1</sup> | Description | Nested in<sup>2</sup> |
| --- | --- | --- | --- |
| $D_{ys}$ | $\lambda, \delta$ | Dysploidy (descending or ascending) | $D_{ys}D_{up}, D_{ys}D_{up}D_{em}^*, D_{ys}D_{up}D_{em}, D_{ys}B_{num}, D_{ys}D_{up}B_{num}$ |
| $D_{ys}D_{up}$ | $\lambda, \delta, \rho$ | Dysploidy and duplication | $D_{ys}D_{up}D_{em}^*, D_{ys}D_{up}D_{em}, D_{ys}D_{up}B_{num}$ |
| $D_{ys}D_{up}D_{em}^*$ | $\lambda, \delta, \rho = \mu$ | Dysploidy, constraining equal rates of duplication and demi-polyploidy | |
| $D_{ys}D_{up}D_{em}$ | $\lambda, \delta, \rho, \mu$ | Dysploidy, duplication, and demi-polyploidy | $D_{ys}D_{up}D_{em}^*$ |
| $D_{ys}B_{num}$ | $\lambda, \delta, \beta, v$ | Dysploidy and base number transition | $D_{ys}D_{up}B_{num}$ |
| $D_{ys}D_{up}B_{num}$ | $\lambda, \delta, \rho, \beta, v$ | Dysploidy, base number transition, and duplication | |

<sup>1</sup> The model parameters are the base number ( $\beta$ ), and rates of ascending dysploidy ( $\lambda$ ), descending dysploidy ( $\delta$ ), duplication ( $\rho$ ), demi-duplication ( $\mu$ ), and base number transition ( $v$ ).
<sup>2</sup> In case all parameters of the model are a subset of other models, the more complex models are indicated.

In a common application of chromEvol, several models are fitted to *D*, the optimal model is selected based on its relative fit using established model selection criteria (e.g. AIC), and subsequent inference tasks are performed based on this model. The model adequacy test can be carried out to any model of interest, whether or not it is the most fitted one. The general aim of this test is to examine whether a specified model, *M*_*x*_, is able to generate data that are similar to *D*. Our model adequacy procedure is based on parametric bootstrapping (Goldman, 1993; Efron & Tibshirani, 1994), where the observed data are compared to a background distribution generated from simulations. These simulations are generated under the specified model, whose parameters, 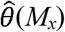, were optimized with respect to *D* and the respective probabilities of chromosome-numbers inferred at the root of the phylogeny (exact details of the simulation procedure are given in the Supporting Information). Comparing between true and simulated data is performed using a set of test statistics, which reflects various characteristics of the data. First, the test statistics (*T*_*1*_, *T*_*2,…,*_*T*_*m*_ ; *m* denotes for the number of statistics) are computed for the true data *D*. Second, multiple datasets are simulated under the specified model and its inferred parameters. For each simulated dataset, the same set of test statistics is computed, resulting in a distribution for each test statistic (*T*_*s1,*_ *T*_*s2*_,…, *T*_*sm*_). If the empirical value of the test statistic falls within the range of variation encompassed by the simulated data (herein defined as the 2.5^th^ and 97.5^th^ percentiles), the model is considered as capable of generating data similar to the original ones and is thus inferred as adequate. Otherwise, it is inferred as inadequate. A schematic illustration of the developed model adequacy test is presented in Fig. **1**.

**Fig 1.**
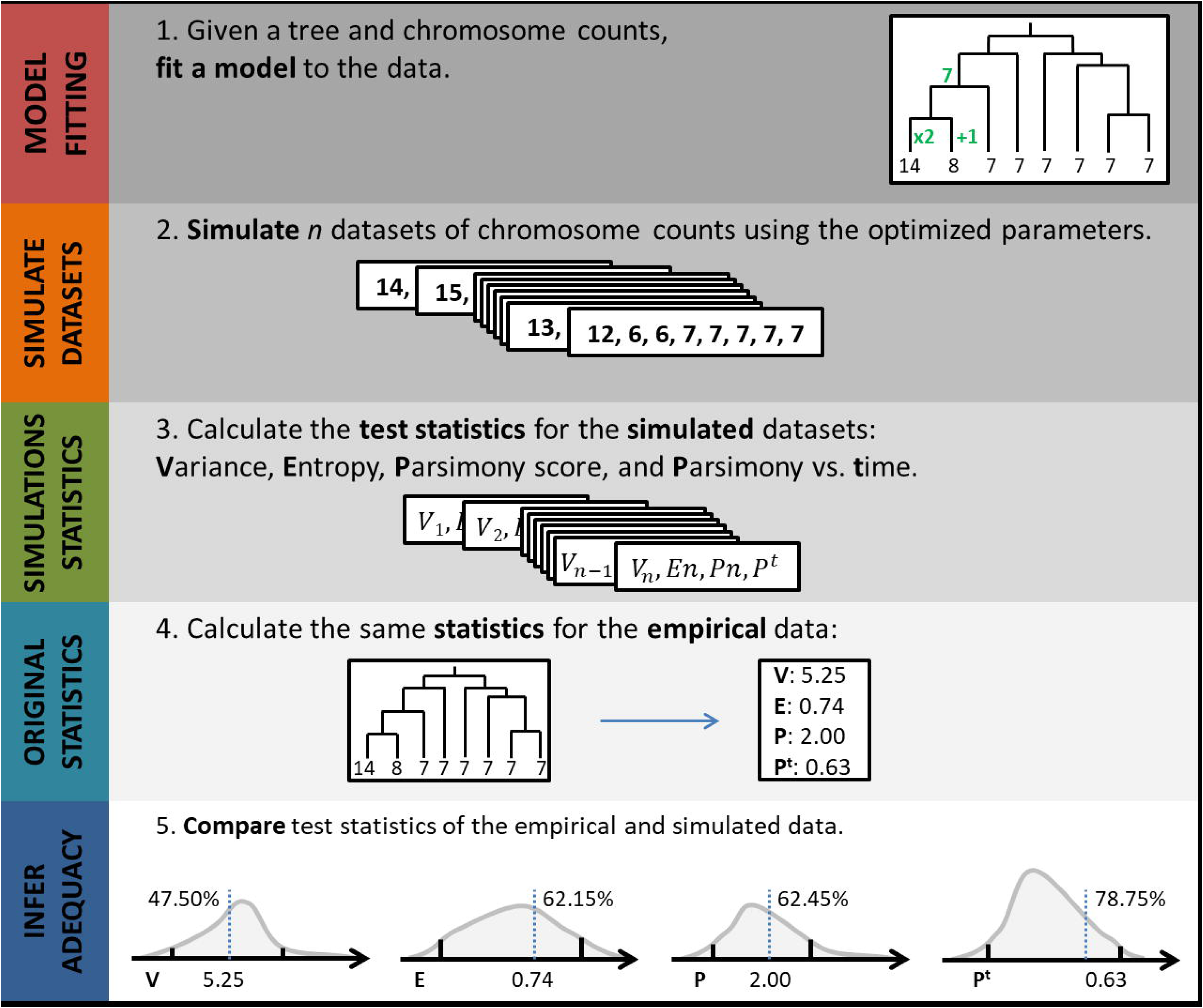
A schematic illustration of the model adequacy framework for likelihood models of chromosome-number evolution. In the case illustrated here, the model is adequate because none of the test statistics lies in the tails of the simulated distribution.

In our implementation, four test statistics were calculated given the chromosome-number data of extant taxa and the corresponding phylogeny: (1) Variance; higher values in the simulated data relative to the observed ones may point to some constraints that were not accounted for by the model (e.g. hard bounds on the number of chromosomes in the genome), or to errors in the parameter estimation process. (2) Shannon’s entropy (Shannon, 1948); Lower entropy of the observed data than predicted by the model is indicative of higher-than-expected concentration of genomes with certain haploid numbers. This could be due to selective constraints, or to a very low variability exhibited in certain subclades of the phylogeny, such that specific states are clumped into large blocks of the tree more than expected. (3) Parsimony score; the most parsimonious number of character transitions across the phylogeny is calculated based on Fitch (1971). If the parsimony score of the observed data are lower than expected it means that the model assumes more transitions than actually occurred. This could occur due to rate heterogeneity across the tree. For example, if chromosome-number transitions occur more frequently in one subclade relative to the rest of the phylogeny, this could be accommodated by inferring higher values of the transition rates. (4) Parsimony versus time (Pars^Time^); the parsimonious number of transitions are computed per branch using the accelerated transformation criterion (ACCTRAN; Farris, 1970). The regression line between the divergence times (computed from the root to the end of the branch) and their parsimony scores is calculated, and the slope of this line is taken as the test statistic. This statistic is similar in spirit to that employed by Pennell *et al*. (2015) for testing the adequacy of models for continuous trait evolution. Under a time-homogenous model, as implemented in chromEvol, we expect no relationship between the divergence times and the number of transitions. Violations of this assumption suggests that transitions are either concentrated around the root or occur more frequently towards the tips. We note that aside from these four statistics, two additional ones were computed (the range and the number of unique counts). These two statistics were found to be highly correlated with the other test statistics (*r*^*2*^ = 0.85 between range and variance and *r*^*2*^ = 0.74 between unique counts and entropy, when computed over the 200 empirical datasets; detailed below), and thus we chose to discard them from further analyses. The coefficient of determinations between all pairs of the four remaining test statistics was below 0.40 (Supporting Information Table S1). Because the four test statistics are not independent and researchers might be interested in revealing the specific aspects of the data that differ from expectations, we followed Pennell *et al*. (2015) and did not apply a multiple testing correction. Thus, in all analyses presented here a model is considered as adequate only if all four statistics fall within the boundaries of the simulated distribution.

### Performance assessment using simulations

Simulations were conducted to examine the performance of the model adequacy procedure. Given an input phylogeny and a set of model parameters, simulated chromosome numbers were generated as previously described in Mayrose *et al*. (2010). As the number of simulation conditions is infinite, we concentrated on eight scenarios that vary in terms of data size (the number of tips in the phylogeny and the observed chromosome-number distribution) and the inferred pattern of chromosome-number change (Table 2). The phylogenies, chromosome counts, and model parameters were taken from empirical datasets previously analyzed using chromEvol (Glick *et al*., 2016; Rice *et al*., 2019), thus representing realistic data characteristics. For each simulation scenario, a total of 100 replicates were generated. Each simulated dataset was then fitted to a set of four models: D_ys_, D_ys_D_up_, D_ys_B_num_, D_ys_D_up_B_num_. For each simulation scenario, one of these models was the generating model (i.e. the model that was used to simulate the data) and three were non-generating models. We note that these models share both common and distinct aspects of the parameter space, such that some, but not all, models are nested within each other (Table 1). Finally, the adequacy of each model to the simulated data was assessed.

**Table 2.** The eight simulation scenarios examined in this study.

| Genus | Number<br>of taxa | Generating<br>model | Model parameters<sup>1</sup> |  |  |  |  |
| --- | --- | --- | --- | --- | --- | --- | --- |
| | | | $\lambda$ | $\delta$ | $\rho$ | $\beta$ | $\nu$ |
| *Aloe* | 120 | D<sub>ys</sub>D<sub>up</sub> | 0 (0) | 0.34 (1) | 2.61 (8) |  |  |
| *Phacelia* | 53 | D<sub>ys</sub>D<sub>up</sub> | 0.20 (2) | 2.33 (21) | 0.67 (6) |  |  |
| *Lupinus* | 77 | D<sub>ys</sub> | 0.85 (7) | 9.53 (76) |  |  |  |
| *Hypochaeris* | 38 | D<sub>ys</sub> | 1.14 (5) | 0.43 (2) |  |  |  |
| *Brassica* | 36 | D<sub>ys</sub>B<sub>num</sub> | 1.24 (11) | 0.70 (6) |  | 8 | 0.55 (5) |
| *Pectis* | 49 | D<sub>ys</sub>B<sub>num</sub> | 0 (0) | 0.40 (2) |  | 12 | 0.55 (3) |
| *Crepis* | 81 | D<sub>ys</sub>D<sub>up</sub>B<sub>num</sub> | 2.41 (19) | 0.99 (8) | 0.26 (2) | 8 | 0.18 (1) |
| *Hordeum* | 36 | D<sub>ys</sub>D<sub>up</sub>B<sub>num</sub> | 0 (0) | 0 (0) | 1.80 (5) | 7 | 1.36 (4) |

<sup>1</sup> In parentheses: average number of simulated events across the tree.

### Inference errors of adequate and inadequate models

The consequences of using an adequate versus inadequate model were evaluated by comparing the errors of four common inference tasks: (1) the chromosome number at the root of the phylogeny; calculated as the deviation from 1.0 of the posterior probability assigned to the true (i.e. simulated) chromosome number at the root. (2) The total number of dysploidy events across the phylogeny; calculated as the relative error between the inferred and simulated number of events: 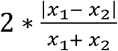, where *x*_1_ and *x*_2_ are the simulated and inferred number of dysploidy events, respectively. In case both *x*_1_ and *x*_2_ equal zero the error was assigned as zero. (3) The total number of polyploidization events across the phylogeny; the relative error was calculated similar to the total number of dysploidy events. Duplication events, demi-duplications, and base-number transitions were regarded as polyploidization events. (4) Ploidy level assignments. The ploidy-level inference of tip taxa, as either diploids or polyploids, was based on the procedure described in Glick & Mayrose (2014). The assignments of all tips were compared between the inferred and true values. The number of falsely inferred taxa, divided by the total number of taxa, was used as the error measure.

In this analysis, six of the eight simulation scenarios in Table 2 were examined. The two scenarios excluded were those generated under the simple D_ys_ model, for which not all inference tasks are relevant. To eliminate possible confounding effects between the specific model used for inference and the magnitude of the error, in this evaluation a single non-generating model (D_ys_D_up_ or D_ys_B_num_) was fitted to the data per simulation scenario (Supporting Information Table S2). For each simulation scenario, 300 replicates were generated. For each replicate, the phylogeny and the simulated chromosome counts were given as input to the model adequacy test and the dataset was determined as either adequate or inadequate. A one-sided *t*-test was conducted to determine whether the error of a certain inference task is significantly larger in the inadequate set compared to the adequate set.

### Application to empirical datasets

To demonstrate the usability of the model adequacy framework, we applied it to a dataset of 200 angiosperm genera, which were randomly selected from a large database consisting of thousands of plant genera, excluding genera with no variations in chromosome numbers as well as those with less than 5 species with both phylogenetic and chromosome-numbers information. The initial database was used, in part or as a whole, in several previous analyses (e.g. Glick *et al*., 2016; Zhan *et al*., 2016; Salman-Minkov *et al*., 2016; Rice *et al*., 2019). From this database we also selected 40 angiosperm genera that each contains at least one allopolyploid species, based on data from Barker *et al*. (2016). Due to overlaps between these two sets, a total of 233 unique datasets were analyzed. Full details of the reconstruction of the original database are described in Rice *et al*. (2019). Briefly, for each genus, the OneTwoTree pipeline (Drori *et al*., 2018) was used to automatically reconstruct the phylogeny using publicly available sequence data as appear in GenBank (Benson *et al*., 2013). Chromosome numbers for all species were retrieved from the Chromosome Counts Database (CCDB; Rice *et al*., 2015). These data were given as input to chromEvol, which was executed on the six models detailed in Table 1. Additionally, we applied similar procedures to seven clades of higher taxonomical ranks, including five families, one subfamily, and one tribe. The evolution of chromosome numbers in these clades using chromEvol was previously examined in several studies (Supporting Information Table S3).

### Implementation and availability

The model adequacy procedure was implemented in Python and R (R Core Team, 2013). The source codes and running instructions are available at https://github.com/MayroseLab/chromEvol_model_adequacy. The obligatory inputs are three files obtained through a chromEvol run of the examined model: the summary results file, the tree with the inferred ancestral reconstruction in a NEWICK format, and the original counts file in FASTA format. The program outputs, for each test statistic examined, its value computed from the empirical data, the percentile in which it falls within the simulated distribution, and the 2.5^th^ and 97.5^th^ percentiles of the simulated distribution as indicative for the upper and lower bounds expected under the modelling assumptions. The model adequacy test is also available for on-line use through the chromEvol web-server (http://chromevol.tau.ac.il/), which is currently in a Beta version.

## Results

In this work we developed a statistical framework for testing the adequacy of likelihood models of chromosome-number evolution. In essence, the method tests whether a specified model is capable of generating data that are similar to the data at hand. If not, the model is considered as providing inadequate description of the data, suggesting that other processes than those modeled have driven the evolution of chromosome numbers along the examined phylogeny. We first evaluated the performance of the model adequacy framework using simulations. We then applied it to a large number of real datasets derived from dozens of angiosperm genera, as well as to seven clades of higher taxonomic ranks, that together vary greatly in their extent of divergence time and patterns of chromosome number variation.

### Framework validation

Simulations were used to validate the developed model adequacy approach. Several simulation scenarios were examined, whose phylogenies and simulated parameters were derived from real data analyses and cover various data characteristics (Table 2). In each scenario, a single model was used to generate the data. Given the simulated data, the generating model and three additional models were fitted to the data, and their adequacies were examined. The four examined models are indicated by the type of transitions they allow for: D_ys_, D_ys_D_up_, D_ys_B_num_, and D_ys_D_up_B_num_ (Table 1). In total, eight different simulation scenarios were examined; two for each type of generating model.

We first examined the type I error rate, i.e. inferring the generating model as inadequate. Our results indicated that when considering a single test statistic independently, the error rate is around the expected value of 0.05 (average = 0.02, across the eight simulation scenarios and four test statistics; Supporting Information Table S4). Combining multiple test statistics together, we consider a model as inadequate if one or more of the statistics fell outside the margins of the simulated distributions (see Methodological Description). Under this definition, the percentage of generating models that were inferred as inadequate varied between 0.04 and 0.13 across the eight simulation scenarios (Table 3). When Bonferroni correction for multiple testing was applied, the type I error rate dropped to an average of 0.008. We note however, that the four test statistics are not independent, violating the assumption of this correction.

**Table 3.** The inadequacy rates of the four tested models in the various simulation scenarios examined (100 simulations per tested model per scenario).

| Simulation scenario | Generating Model | Tested Models<sup>1</sup> |  |  |  |
| --- | --- | --- | --- | --- | --- |
| | | $D_{ys}D_{up}$ | $D_{ys}$ | $D_{ys}B_{num}$ | $D_{ys}D_{up}B_{num}$ |
| Aloe | $D_{ys}D_{up}$ | 0.07 | 1.00 | 0.06 | 0.08 |
| Phacelia | $D_{ys}D_{up}$ | 0.04 | 0.99 | 0.04 | 0.03 |
| Lupinus | $D_{ys}$ | 0.06 | 0.08 | 0.06 | 0.07 |
| Hypochaeris | $D_{ys}$ | 0.03 | 0.06 | 0.04 | 0.05 |
| Brassica | $D_{ys}B_{num}$ | 0.14 | 0.98 | 0.05 | 0.06 |
| Pectis | $D_{ys}B_{num}$ | 0.86 | 1.00 | 0.12 | 0.07 |
| Crepis | $D_{ys}D_{up}B_{num}$ | 0.27 | 0.95 | 0.11 | 0.08 |
| Hordeum | $D_{ys}D_{up}B_{num}$ | 0.77 | 1.00 | 0.30 | 0.13 |

<sup>1</sup> The diagonal (white cells) are cases where the generating model is also the tested model. Dark grey represents over-parametrized models, light grey under-parametrized models, and patterned cells miss-parametrized models.

We next examined the capability of the adequacy test to detect models that deviate from that of the generating models. Three types of model misspecification were examined: over-parameterization, under-parameterization, and miss-parameterization. In the case of over-parameterization, the tested model allows for additional types of chromosome-number change (as represented by extra free parameters) than those used to generate the data. This corresponds to cases where the generating model is nested within the tested model (e.g. the generating model is D_ys_D_up_ while the tested model is D_ys_D_up_B_num_). Our results indicated that the performances of over-parameterized models are very similar to that of the generating models (Table 3). The few discrepancies were the result of either (1) inaccurate parameter estimates of the more general model due to the extra degrees of freedom; (2) the optimization procedure reaching suboptimal regions of the parameter space (we note that while chromEvol allow for more thorough likelihood optimization search, which should reduce such instances, this was not attempted here due to the large number of simulations employed); (3) very similar parameter estimates obtained using the two models, but slight deviations of the test statistics led one model to be inferred as inadequate while the other one as adequate.

In the case of under-parameterized models, the tested model allows for fewer types of transitions than the generating model (e.g. the generating model is D_ys_D_up_ while the tested model is D_ys_). As may be expected, in all simulation scenarios the under-parameterized models were more frequently inferred as inadequate compared to the generating models. The adequacy rate was very low when the tested model allowed only for dysploid transitions while in reality polyploid transitions (either WGD and/or base-number transitions) have occurred (Table 3; all cases where the tested model is D_ys_). The adequacy rates were higher when the generating model allowed for multiple types of polyploid transitions (i.e. D_ys_D_up_B_num_ allowing for both exact duplications and base-number transitions), while the tested model allowed for a subset of these (D_ys_D_up_ and D_ys_B_num_ that allow only for duplications or base-number transitions, respectively). Comparing the adequacy of the two under-parametrized models (D_ys_D_up_ and D_ys_B_num_), the D_ys_B_num_ model that incorporated base-number transitions had higher adequacy rates compared to the D_ys_D_up_ model that allowed for exact duplications, as the former allows for several transitions that frequently include also exact duplications (e.g. in case the base number is 8, both 8→16 and 8→24 transitions are allowed).

In the case of miss-parametrization, the tested and generated models are not nested within each other and thus their parameters only partially overlap. For the set of models examined here, this fits the case where the generating model is D_ys_D_up_ while the tested model is D_ys_B_num_, or vice versa. When the tested model was D_ys_B_num_, it obtained similar adequacy rates to those of the generating D_ys_D_up_ model. In contrast, and similar to the results detailed in the case of under-parameterized models, the D_ys_D_up_ model was inferred as inadequate a large number of times when the generating model was D_ys_B_num_.

### Inference errors of adequate and inadequate models

A central usage of probabilistic models of chromosome number evolution is their inference capabilities, such as ancestral reconstructions of chromosome numbers, or predicting the branches in which dysploidy and polyploidy events have most likely occurred. Still, it is unclear whether the use of inadequate models would deteriorate the performance of such inference tasks. To this end, simulations were used to compare the errors of the following four common inference tasks when adequate and inadequate models are employed: (1) the chromosome number at the root of the phylogeny; (2) the total number of inferred dysploidy events; (3) the total number of inferred polyploidization events, and (4) inferring the ploidy level of tip taxa as either diploid or polyploidy (see Methodological Description for details regarding the error computed for each inference task).

Our results demonstrated that the use of inadequate models frequently leads to larger inference errors, although under some simulation scenarios the inference errors of inadequate models were similar to that obtained using adequate models. For example, the error in the inference of the root chromosome number was significantly larger in the case of inadequate models under two simulation scenarios, but was non-significantly different in the other four (Fig. **2**). Similarly, in two out of the six simulation scenarios, the error of inferring the ploidy level of extant taxa was significantly larger when computed using inadequate versus adequate models. In this case, the magnitude of the error was relatively low whether adequate or inadequate models were applied: when inadequate models were applied, the mean error was 4.6% across all simulation scenarios, reaching up to 12% under the *Brassica* simulation scenario. In comparison, the mean error was 2% when adequate models were applied, reaching up to 6% of erroneous inferences under the *Hordeum* simulation scenario. Larger differences in the errors between adequate and inadequate models were observed in inferring the total number of polyploidizations, and even more so in inferring the total number of dysploidy events. For both these inference tasks, significant differences between adequate and inadequate models were obtained for three out of the six simulation scenarios. Generally, the relative error in inferring the total number of dysploidy events was larger compared to that of inferring the total number of polyploidizations (the mean relative error was roughly twice for dysploidy compared to polyploidy transitions, both in the adequate set and the inadequate set; Fig. **2**).

**Fig. 2.**
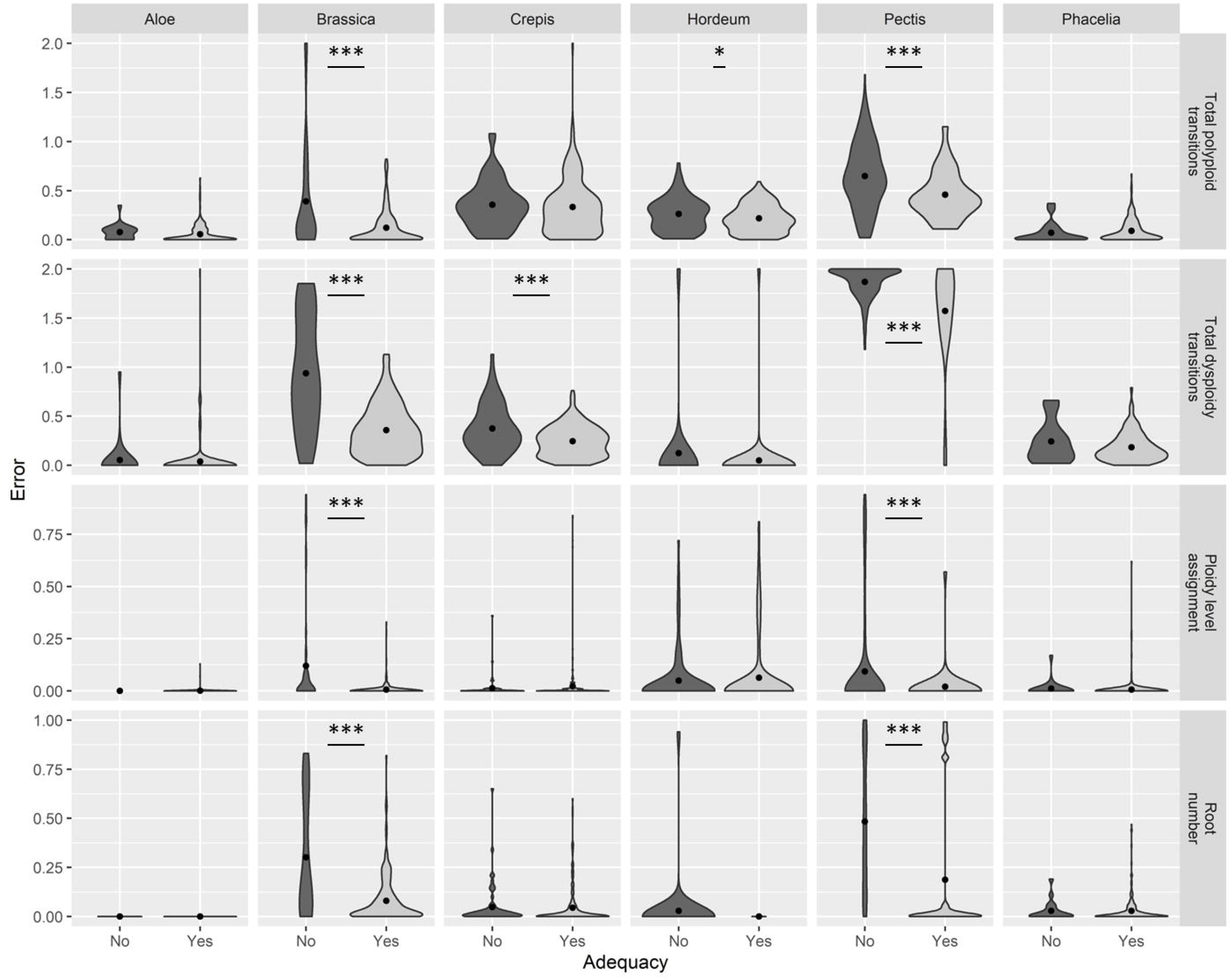
The mean inference errors obtained under adequate and inadequate models for each simulated scenario. Each row presents the error of a different inference task. From top to bottom: inferring the total number of polyploid events across the tree, inferring the total number of dysploid events across the tree, ploidy level assignments of extant taxa, the probability of the chromosome number at the root of the phylogeny. Each column denotes a different simulation scenario. For each scenario, 300 simulations were conducted and runs were partitioned to adequate and inadequate models. The violin plots represent the distribution of the errors obtained for the adequate (light grey, right) and inadequate (dark grey, left) sets. The black dot within each distribution denotes its mean. Asterisk indicates significant difference between the two groups (*, *p* < 0.05 and ***, *p* < 0.01).

### Application to empirical datasets

We applied the model adequacy framework to 200 datasets, each corresponding to a single randomly-selected angiosperm genus. First, we performed a standard model selection procedure based on the AIC (Akaike, 1974) to evaluate the relative fit of each of the six chromEvol models to the data. In 24% of the datasets, the simple D_ys_ model, which allows for dysploid transitions only, was selected. The model that was most frequently selected was D_ys_D_up_ (28%), while models that allow for demi-polyploidy transitions and those that allow for base-number transitions were selected in 27% and 21% of the datasets, respectively (Fig. **3a**). Next, we applied the model adequacy test to the best model identified for each dataset. We found that in 74% of the genera, the model that was chosen as best by the AIC was inferred to provide an adequate description of the data. Applying the model adequacy test to all six models per dataset (whether or not selected as best), we found that models that allow for fewer types of transitions were more frequently predicted as inadequate (Fig. **3b**). For example, the D_ys_ model that allows only for dysploidy transitions was adequate in only 28% of the 200 datasets, models that additionally allow for one type of polyploidy, either duplication or base-number transition, were adequate 60% and 64% of the cases, respectively, while the three models that incorporate two types of polyploidy transitions (D_ys_D_up_D_em_, D_ys_D_up_D_em_ *, and D_ys_D_up_B_num_) were inferred as adequate most frequently. The adequacy rates of all models were generally related to the complexity of the model that was selected as optimal. Thus, when the most complex models were selected (D_ys_D_up_D_em_ and D_ys_D_up_B_num_), the adequacy rates of all models – including that of the chosen model – were low (33% and 47%, respectively), while when the least complex model was selected, the adequacy rates of all models was high (70%; Supporting Information Table S5).

**Fig. 3.**
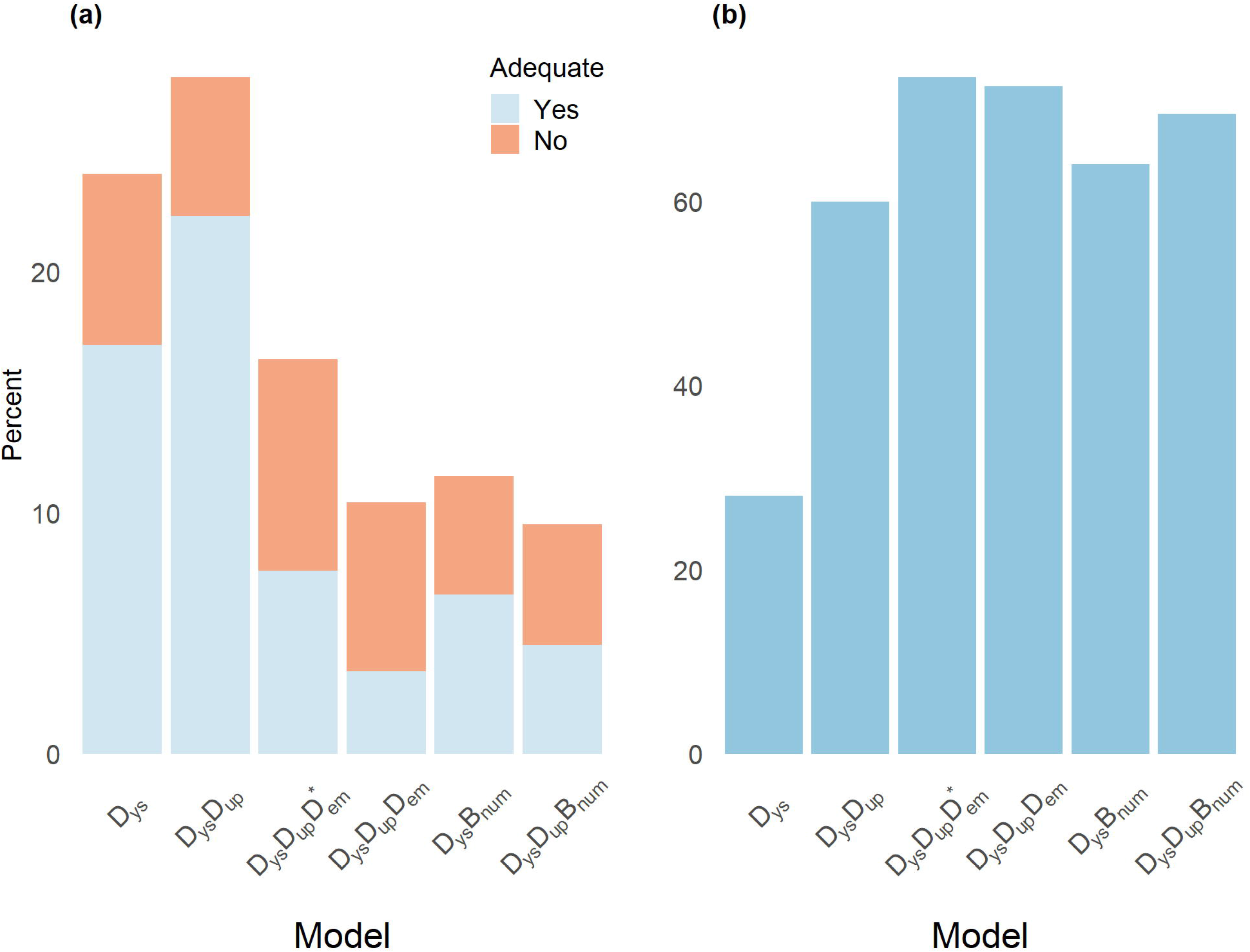
Application of the model adequacy test to 200 angiosperm genera. (**a**) A bar plot representing the frequency of selection according to the AIC of each of the six tested models in the 200 examined angiosperm genera. The height of each bar is partitioned according to the percentage of genera that were determined as adequate (light blue) or inadequate (red). (**b**) The adequacy rate of each model when applied to all genera, regardless of whether the model was selected (*n =* 200).

Next, we examined the model adequacy procedure in groups that have evolved via reticulate evolution at some point in their histories. In these clades, the underlying assumption of the chromEvol framework, in which evolution proceeds along a phylogenetic structure, is violated, at least to some extent. This analysis was performed on 40 genera that were identified in the literature to include allopolyploid species, and thus hybridizations were reported to occur (data taken from Barker *et al*., 2016). In the majority of these genera (24 out of 40), the model that was selected as optimal according to the AIC was found by our model adequacy procedure as inadequate. This adequacy rate is significantly lower (*p* ≪ 0.05; *χ*^2^ test) compared to a random set of 193 genera in which allopolyploidy was not reported (the 200 genera analyzed above, omitting seven that include a reported allopolyploid species).

Finally, we evaluated the model adequacy procedure on a set of seven groups whose taxonomic rank is higher than the genus level, thus representing clades whose divergence time is generally older than those inspected above. The evolution of chromosome numbers in these clades likely violates the time homogeneity assumption of chromEvol, in which the transition pattern is similar across the phylogeny. For four of these seven clades, the model that was chosen as optimal according to AIC did not provide adequate description of the data according to the model adequacy test (Supporting Information Table S3) and in one additional case the empirical values of two test statistics were placed close to the lower boundaries of the simulated distributions (falling in the 0.027 and 0.043 percentiles). Taken together, the last two analyses indicate that the model adequacy procedure can identify cases in which the evolution of chromosome numbers is driven by processes that deviate from the basic modelling assumptions of the chromEvol framework.

## Discussion

For over a century, the determination of chromosome numbers has played a vital role in studying evolutionary and genomic processes in plants. Probabilistic models of chromosome-number change are a relatively recent addition to the research toolbox available to study the evolution of major genomic processes. As the usage of such models increases, so does the need to assess their validity when applied to real data. Here, we developed a model adequacy test for likelihood models of chromosome-number evolution. We focused our analysis on those models implemented in the chromEvol software (Glick & Mayrose, 2014), but the procedures are general and can be implemented in other platforms that use variations to the chromEvol model (Freyman & Höhna, 2017; Zenil-Ferguson *et al*., 2017; Blackmon *et al*., 2019). The developed test is based on the parametric bootstrapping approach (Goldman, 1993; Efron & Tibshirani, 1994) in which observed data are compared to a simulated distribution generated by the examined model. Using multiple test statistics that describe various characteristics of the data, the test allows to determine whether the model can generate data that are similar to those found in the observed ones.

Our simulation results indicate that the model adequacy framework has an acceptable type I error rate (i.e. inferring as inadequate a model that was used to generate the data). However, higher type I errors were found in models that allow for base-number transitions (D_ys_B_num_ and D_ys_B_num_D_up_). This suggests that these models might not be appropriate in all cases. The current implementation of such models assumes the same rate for all possible base-number transitions (e.g. given a base number of *β* = 7, the additions of 7, 14, or 21 chromosomes are equally likely). Alternatively, it may be more appropriate to place a probability distribution over the possible base-number transitions. This will allow, for the example of *β* = 7, higher rates for additions by 7 chromosomes compared to those by 21.

Our simulation results also demonstrate that the adequacy rate of over-parameterized models, which allows for more types of transitions than those that truly occurred, is similar to that of the generating models. While it is expected that the accuracy of inferring the model parameters will decrease as overly-complexed models are evaluated, in many cases the auxiliary parameters were optimized to very low values, resulting in a process that is nearly identical to the generating model. Thus, it seems that the flexibility offered by complex models does not necessarily lead to their disadvantage, at least for some inference tasks, as has been recently demonstrated for models of nucleotide sequence evolution (Abadi *et al*., 2019). In other cases of model violations, either for under-parameterized or miss-parameterized models, when the rate parameters deviated substantially from the original ones (e.g. dysploidy rates an order of magnitude larger than the simulated rates), the model adequacy framework detected such cases as inadequate. This suggests that the adequacy test is capable of detecting models that are completely wrong. In other cases, the nature of model misspecification affected the outcome. In the simulations examined here, D_ys_B_num_ was more frequently adequate than D_ys_D_up_, both in the case of under-parameterization (i.e. when the generating model was D_ys_D_up_B_num_ such that both models miss one type of transition) and miss-parameterization. Nevertheless, we note that the D_ys_B_num_ model may not fit well in large phylogenies with high dysploidy rates. In its current implementation, the model assumes that a single base number typifies a clade. However, if there is a high dysploidy rate, each subclade of the phylogeny may be characterized by its own base number or by multiple base numbers, which will necessitate more complex modelling options.

We further tested the consequences of using an inadequate model by examining the errors of several inference tasks. First, we found that the difference in inference error between adequate and inadequate models depended on the simulation scenarios: in some simulation scenarios the use of inadequate models resulted in significantly inflated inference errors compared to the use of adequate models, in some scenarios it affected only certain inference tasks and not others, while in others the difference was negligible for all tasks. Second, we found that some inference tasks are much more sensitive to model misspecification than others. The assignment of extant taxa as diploids or polyploids was the inference task that was least affected from using an inadequate model, and in general, the error of this inference task was very low (in all scenarios, the ploidy level of 88% or more of the taxa were correctly identified). This indicates that determining the ploidy levels of extant taxa is generally robust to model misspecification. On the other hand, the error of determining the number of events that had occurred – either dysploid or polyploid transitions – can be substantial when inadequate models are employed.

Applying the model adequacy test to hundreds of angiosperm genera, we found that in the majority of the cases the best-fitted model provided sufficient approximation to the evolutionary processes underlying the data and was determined as adequate. However, in roughly one fourth of the examined genera, this selection turned out to be inadequate, suggesting that there is ample room for future modelling improvements. Indeed, we found high rates of model inadequacy when applying the developed procedures to two types of clades that are expected to violate basic modelling assumptions: first, clades in which allopolyploidy events are known to occur, thus violating the assumption that evolution proceeds via a phylogenetic structure; second, in the case of large and diverse clades in which a single transition process is fitted to the entire phylogeny, following the time homogeneity assumption, is insufficient. These results thus indicate that promising future developments would be to focus on analytical procedures based on phylogenetic networks (Nakhleh, 2010), rather than on bifurcating phylogenies, and to further incorporate time-heterogeneous processes.

Phylogenetic model adequacy tests have been previously developed for other data types and inference tasks, although their use has not been widely adopted. This could be due to the apparent limited benefit offered to a researcher when all examined models are deemed inadequate when applied to a clade of interest. We argue, however, that model adequacy tests are of practical use to methods developers and end users alike, and should thus be regularly practiced as part of a broader model assessment routine. For researchers interested in data analysis, inadequate models can hint on errors in the input data, which should thus be more carefully inspected. In the case studied here, possible sources of errors include those in the assumed phylogenetic hypothesis, in the collection of chromosome counts, or in taxa sampling. Inadequacy could also point to additional attributes that should be considered in the analysis. For example, if all models that assume a time-homogenous transition process fail, it could suggest that patterns of chromosome-number change are dependent on an organismal trait (e.g. the plant growth form), that if accounted for, using more complex models (e.g. Zenil-Ferguson *et al*., 2017; Blackmon *et al*., 2019) would enhance the analysis. For researchers interested in large scale analyses that include multiple datasets, where the in-depth examination of each inadequate dataset is not feasible, the filtration of such clades is one obvious possible direction. For some inference tasks, such as the identification of ploidy levels of extant taxa, the effect of using an inadequate model is rather negligible, indicating that the treatment of the flagged clades should be tuned to the analysis in question. For the developers, the frequent application of model adequacy tests should provide interesting test cases on which new models are trained. Moreover, when a model is deemed inadequate, the test statistics that fail to align may point to processes absent from existing models, which could be included in the future. Model adequacy should thus take a vital part in this recurrent chain of scientific progress in which new methods are developed, regularly used, and then replaced by more advanced alternatives.

## Acknowledgements

A.R. is supported by a fellowship from the Edmond J. Safra Center for Bioinformatics at Tel Aviv University and by the NA’AMAT Professional Scholarship. This study was supported by grant # 961/17 from the Israel Science Foundation to I.M.

## Author Contribution

IM and AR conceived the study; AR built the tool and analyzed the data; AR and IM wrote the manuscript; IM supervised the study.

## Supplementary information

### Supplementary Information Methods

Item 1: Description of the simulation procedures.

### Supplementary Information Tables

Table S1: Pearson’s *r* coefficient between each pair of statistics.

Table S2: The generating and fitted model for each simulation scenario used in the comparison of inference error between adequate and inadequate models.

Table S3: Details of the seven plant clades, whose taxonomic rank is above the genus level, examined in this study.

Table S4: Type I error rates for each test statistic per simulation scenario.

Table S5: Adequacy rates of all models, including those of the chosen models.

