## Supporting Information for "Model Adequacy Tests for Likelihood Models of Chromosome-Number Evolution"

**Supplementary Information Methods**

#### Model adequacy simulations

Simulations constitute a central step in the model adequacy procedure as they are used to generate artificial data that correspond to the model assumptions and whose parameters were optimized to best fit the examined empirical data. In our case, the simulations generate chromosome numbers for extant taxa, represented by the tips of the input phylogeny. For each dataset, *n* simulations are performed (by default, *n* = 1,000), thus creating a distribution for each of the computed test statistics. Simulated data are prepared by modeling the evolutionary process along the input phylogeny and given the inferred set of model parameters. These simulations are performed using the embedded transition rate matrix of the Markov process, *Q* and following the procedures described in Mayrose *et al.* (2010). Here we describe several details that are needed for the model adequacy procedures.

First, a parameter, *C*, that specifies the upper bound for the number of chromosomes in a genome has to be provided as input for the simulations. This number specifies the size of the simulated rate matrix and thus influences the running time of generating the simulated data. When *C* is very large (i.e., over 1,000), the simulations can take prohibitively long time. On the other hand, if this value is too low, it may be reached during the simulations, thus rendering the simulations inappropriate (i.e., once the process has reached this number it cannot proceed to higher values). The following iterative procedure was thus employed to set *C* to an appropriate value:
(1) The maximum chromosome number in the empirical data is found ($x_{max}$).
(2) The initial value of *C* is set $C=\max\left\{ 200,{10+x}_{max} \right\}$
(3) *n* simulations are performed with the current value of *C*.
(4) If the upper bound is reached in one of the simulations, the value of *C* is increased to C+100 and step (3) is repeated. Otherwise, the obtained *n* simulated datasets are used for the model adequacy test.

Second, a special treatment was given to simulations that allow for base-number transitions (e.g. when using the D_ys_B_num_ and D_ys_D_up_B_num_ models; Table 1, main text). All transition types, aside from base-number transition, specify a single event while being in a genome with *i* haploid chromosomes. For example, ascending dysploidy specifies the transition *i*🡪*i*+1 with rate *λ,* while WGD specifies the transition *i*🡪2*i* occurring at rate *ρ*. In contrast, the incorporation of base-number transitions allows for several possible moves: *i*🡪*i*+*β*, *i*🡪*i*+2*β,* *i*🡪*i*+3*β,* etc, each occurring at rate *ν*. Thus, the size of the rate matrix used to simulate the data affects the number of entries with positive values and could thus affect the number of simulated transitions. As noted above, however, the size of the transition matrix used to simulate the data could be of different size than that used for fitting the model to the empirical data, meaning that the two models are incongruent. We thus included an additional parameter, *U*, that specifies the maximal range of base-number transitions allowed and also affect the value of the inferred *ν* parameter. For example, if this upper range is 36 and *β* = 12 then three moves are allowed away from the current state (*i*🡪*i*+12, *i*🡪*i*+24, *i*🡪*i*+*36*). The incorporation of this parameter also allows the model to focus on a more succinct set of realistic moves. To fix this value to realistic values, when base-number transitions are allowed, the simulations are performed in a two-step manner. First, fitting the model to the empirical data, while setting *U* to the range (maximum – minimum) of chromosome-numbers found in the data. Second, we used the inferred ancestral states of this initial run to obtain an estimate of the chromosome-number transitions that have occurred along all branches of the phylogeny. In case the largest transition found is smaller than the initial value of *U*, another round of optimization is performed, while setting *U* as the largest transition.

### Supporting Tables

**Table S1.** Pearson’s *r* coefficient between each pair of statistics, calculated from the statistics percentiles in the set of angiosperm genera tested in this study (*n* = 200).

|  | Variance | Entropy | Parsimony | Pars^Time^ |
| --- | --- | --- | --- | --- |
| Variance |  | 0.53 | 0.10 | -0.33 |
| Entropy |  |  | 0.63 | -0.23 |
| Parsimony |  |  |  | 0.22 |
| Pars^Time^ |  |  |  |  |

**Table S2.** The generating and fitted model for each simulation scenario used in the comparison of inference error between adequate and inadequate models.

| Simulation scenario | Generating model | Fitted model | Adequacy (%)^1^ |
| --- | --- | --- | --- |
| *Aloe* | D_ys_D_up_ | D_ys_B_num_ | 92.67 |
| *Phacelia* | D_ys_D_up_ | D_ys_B_num_ | 93.67 |
| *Brassica* | D_ys_B_num_ | D_ys_D_up_ | 82.67 |
| *Pectis* | D_ys_B_num_ | D_ys_D_up_ | 11.67 |
| *Crepis* | D_ys_B_num_D_up_ | D_ys_D_up_ | 79.00 |
| *Hordeum* | D_ys_B_num_D_up_ | D_ys_B_num_ | 67.67 |

^1^The adequacy percentage is computed as the number of simulations in which the fitted model was determined as adequate across 300 replicates per simulation scenario.

**Table S3.** The seven plant clades, whose taxonomic rank is above the genus, examined in this study. The best model is the one chosen according to the AIC. Clades marked in bold are those in which the best model was determined as adequate. Each diagram under the ‘Test statistic’ column represents the percentiles of the four empirical test statistics in the simulated distribution: red arrow denotes for a test statistic that falls in the tails of the distribution (< 0.025 or > 0.975) and green arrow denotes a test statistic that falls within the central 95 percentiles of the distribution.

| Clade | Best model | Range of counts (unique counts) | Number of tips (genera) on tree | Test statistics |
| --- | --- | --- | --- | --- |
| Antirrhineae  (Ogutcen & Vamosi, 2016) | D_ys_D_up_B_num_ | 22 (12) | 124 (19) |  |
| Aspleniaceae  (Schneider *et al.*, 2017) | D_ys_D_up_D_em_ | 252 (10) | 94 (3) |  |
| Bombacoideae  (Costa *et al.*, 2017) | D_ys_D_up_D_em_^*^ | 37 (14) | 129 (8) |  |
| **Cistaceae  (Aparicio *et al.*, 2019)** | **D_ys_D_up_** | **13 (7)** | **70 (8)** |  |
| **Melanthiaceae  (Pellicer *et al.*, 2014)** | **D_ys_D_up_B_num_** | **27 (12)** | **108 (16)** |  |
| Passifloraceae  (Sader *et al.*, 2019) | D_ys_D_up_D_em_^*^ | 37 (14) | 129 (8) |  |
| **Pontederiaceae  (Ness *et al.*, 2011)** | **D_ys_D_up_** | **19 (6)** | **15 (4)** |  |

**Table S4.** Type I error rates for each test statistic per simulation scenario. The different scenarios are denoted by the genus name whose tree and inferred model parameters were used to simulate the data. For each scenario, 100 simulations were conducted.

| Simulation scenario | Test statistic | | | |
| --- | --- | --- | --- | --- |
|  | Variance | Entropy | Parsimony | Pars^Time^ |
| *Aloe* | 0.03 | 0.01 | 0.00 | 0.04 |
| *Phacelia* | 0.01 | 0.00 | 0.00 | 0.03 |
| *Lupinus* | 0.01 | 0.02 | 0.01 | 0.05 |
| *Hypochaeris* | 0.01 | 0.01 | 0.00 | 0.04 |
| *Brassica* | 0.00 | 0.00 | 0.00 | 0.05 |
| *Pectis* | 0.04 | 0.02 | 0.02 | 0.07 |
| *Crepis* | 0.02 | 0.03 | 0.00 | 0.05 |
| *Horderum* | 0.11 | 0.01 | 0.00 | 0.03 |

**Table S5.** Adequacy rates of all models, including those of the chosen models.

| Best model | Number of selections^1^ | Total adequacy rates^2^ (%) |
| --- | --- | --- |
| D_ys_ | 48 | 70.14 |
| D_ys_D_up_ | 56 | 79.47 |
| D_ys_D_up_D_em_^*^ | 33 | 46.43 |
| D_ys_D_up_D_em_ | 21 | 32.80 |
| D_ys_B_num_ | 23 | 57.24 |
| D_ys_D_up_B_num_ | 19 | 47.37 |

^1^ Number of times the model was selected as best model out of the 200 tested cases.

^2^ Calculated over the whole dataset when all six models were fitted with the respective model being chosen as best model.
